# Neuregulin-1 protects against respiratory viral induced mortality

**DOI:** 10.1101/2023.05.10.540232

**Authors:** Syed-Rehan A Hussain, Michelle Rohlfing, Jennifer Santoro, Jenny Resiliac, Phylip Chen, Mark E. Peeples, Mitchell H Grayson

## Abstract

Respiratory viral infections due to RNA viruses such as respiratory syncytial virus (RSV) and influenza lead to significant morbidity and mortality. Using a natural rodent pathogen similar to RSV, Sendai virus (SeV), we found that mice made atopic with house dust mite before viral infection all survived a normally lethal SeV infection.

Moreover, adoptive transfer of CD11c^+^ cells from atopic mice delayed viral mortality. Neuregulin-1 (NRG1) message was highly expressed in CD11c^+^ cells from atopic mice and atopic lungs and bronchoalveolar lavage fluid had elevated levels of NRG1 protein. Administration of NRG1 protected non-atopic mice from death and associated with reduced alveolar epithelium permeability. Utilizing an *in vitro* system of well-differentiated human bronchial epithelial cells and mouse tracheal epithelial cells NRG1 reduced RSV and SeV titers. Expression of genes that play a role in airway epithelium integrity and stability were altered by NRG1; potentially regulating viral induced dysregulation of the epithelia and suggesting NRG1 mediated maintenance of homeostasis. In conclusion, our studies demonstrate atopy induced NRG1 likely plays a novel role in survival from severe respiratory viral infections and may have therapeutic value to prevent mortality from these infections.

**Significance:** Severe respiratory viral infections are associated with significant mortality in infants and the elderly; however, allergic disease can protect from these outcomes. This study identified a protein called neuregulin-1 (NRG1), produced by cells of the immune system in allergic mice, that provides a survival advantage against respiratory viral infection. NRG1 pretreatment in non-atopic mice infected with a lethal dose of a rodent RNA virus (Sendai virus), similar to human respiratory syncytial virus, significantly reduced death. Further, NRG1 pretreatment reduced viral replication in human and mouse airway epithelial cell cultures. These studies signify a potential therapeutic role of NRG1 in modulating the severity of respiratory viral infections.

## Introduction

Respiratory viral infections with negative-strand RNA viruses, such as influenza virus (IAV), respiratory syncytial virus (RSV) and parainfluenza virus (PIV) are a major cause of morbidity and mortality. Several groups have demonstrated pre-existing atopy protects from mortality to IAV in mouse models (Ishikawa et al., 2012; An et al., 2018; Furuya et al., 2015). Human data supports this relationship – in the 2009 H1N1 IAV pandemic patients with asthma hospitalized with IAV had less severe outcomes, while a recent meta-analysis concluded patients with asthma had a significantly lower mortality to COVID-19, and the presence of food allergies was reported to have reduced the likelihood of being infected with SARS-CoV-2 (Veerapandian et al., 2018; Louie et al., 2009; Myles et al., 2012; Hou et al., 2021; Seibold et al., 2022). The mechanistic explanation for why pre-existing atopy would be protective is not known, although various explanations have been provided for the increased survival in animal models including activation of NK cells, induction of Type III interferon and expression of TGFβ (Ishikawa et al., 2012; An et al., 2018; Furuya et al., 2015).

RSV is a major viral pathogen, especially for infants and the elderly. In the United States annually there are on average 2.1 million outpatient visits and 58,000 hospitalizations for RSV in children under 5 years of age. (Hall et al., 2009; Rha et al., 2020). Globally over 200,000 children under 5 years of age die annually of RSV infections (Li et al., 2022). In those 65 years of age or older, RSV accounts for an average of 177,000 hospitalizations and 14,000 deaths annually, with similar mortality rates to influenza (IAV) (Chorazka et al., 2021; Falsey et al., 2005). Reducing severity of an RSV infection has potential for significant clinical impact by reducing mortality in infants and adults. However, it is not known if pre-existing atopy has similar protection against RSV infection as seen with IAV.

We have a high-fidelity model of respiratory viral infection. Our mouse model utilizes a natural rodent pathogen, Sendai virus (SeV), which is closely related to RSV, but unlike RSV, faithfully replicates in mouse airway epithelial cells (Hussain et al., 2021; Cheung et al., 2010a; Grayson et al., 2007a; Grayson et al., 2007b). Infection with SeV leads to acute bronchiolitis followed by a chronic inflammatory response associated with airway hyper-reactivity and mucous cell metaplasia. These are the cardinal features of human paramyxoviral (PIV), pneumoviral (RSV), and orthomyxoviral (IAV) infection, and we have used this model to identify a novel immune axis, which we have validated in humans (Khan and Grayson, 2010; Sammon et al., 2019).

Using this model, we recently defined a mechanistic pathway that explains how pre-existing atopy could prevent development of post-viral airway disease (Hussain et al., 2021). During this study we noted that SeV infected atopic mice lost less weight than non-atopic mice, suggesting a possible survival benefit of atopy. In the current report we demonstrate that atopy protects from SeV induced mortality similar to previous IAV studies (Ishikawa et al., 2012; An et al., 2018; Furuya et al., 2015). Unlike the IAV studies, we now demonstrate that atopic mice have increased lung and airway levels of neuregulin 1 (NRG1), a cytokine of the epidermal growth factor family, that interacts with ErbB receptor tyrosine kinases (Britsch, 2007). Importantly, exogenously administered NRG1 in non-atopic mice significantly reduced mortality to the viral insult. Finally, we demonstrate that *in vitro* NRG1 treatment can reduce SeV replication in well-differentiated mouse tracheal epithelial cells and, more importantly, RSV replication in human bronchial epithelial cells. Our studies demonstrate that NRG1 protects from a lethal respiratory viral infection and may be a mechanism by which pre-existing atopy prevents mortality from respiratory viral infections.

## Results & Discussion

### Pre-existing atopy prevents SeV induced mortality

To model the impact of the atopic state in preventing severe disease from respiratory viral infection, we made C57BL6 mice atopic by sensitizing and challenging mice with house dust mite antigen (HDM (atopic)) or PBS (non-atopic control; NA). We previously demonstrated that inoculating i.n. with 2×10^5^ pfu (“regular” dose) SeV 3d after the last HDM challenge attenuated weight loss, reduced SeV titer, and inhibited SeV induced airway hyperreactivity. This prevention of post-viral disease required the presence of PMN in the atopic mice (Hussain et al., 2021). Given that atopic mice had suppressed respiratory viral induced disease with the regular dose of SeV, we examined the impact of pre-existing atopy on a normally lethal dose of SeV (2×10^6^ pfu; “high” dose).

Interestingly, like with IAV infection in mouse models (Ishikawa et al., 2012; An et al., 2018; Furuya et al., 2015) all mice made atopic before SeV infection survived high dose viral inoculation, while all NA mice did not (Figure 1A). Survival in atopic mice was associated with a significant reduction in viral titer (Figure 1B).

**Figure 1.**
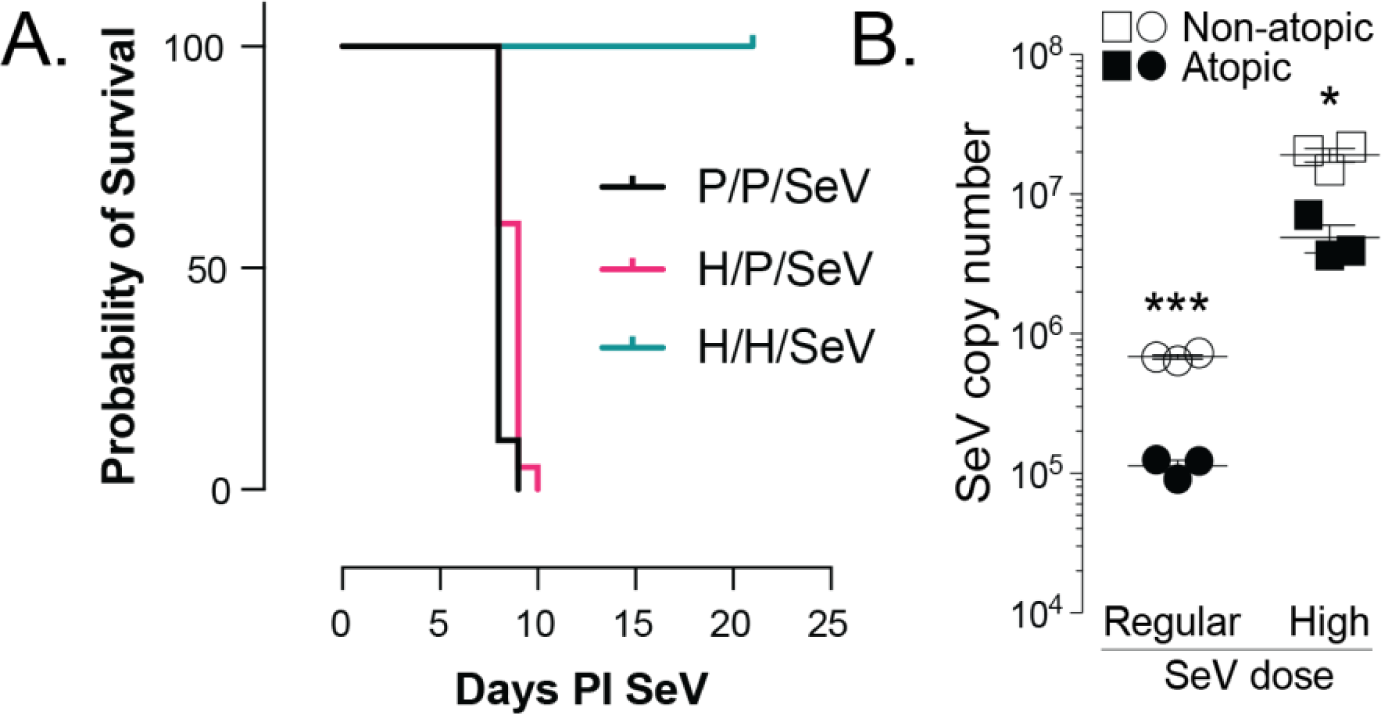
Atopy prevents lethal SeV infection. (**A**) Mice sensitized and challenged with house dust mite extract (“H/H/SeV”, atopic) are protected from mortality to high dose (2×10^6^ pfu) SeV infection, while NA mice (i.e., only sensitized (“H/P/SeV”) or neither sensitized or challenged (“P/P/SeV”)) all succumbed to the viral insult (*p*<0.0001 H/H/SeV versus each of the other NA groups; p=0.016 P/P/SeV vs H/P/SeV; Mantel-Cox; P/P/SeV n= 9; H/P/SeV and H/H/SeV n= 20 combined from three separate experiments). PI = post inoculation. (**B**) Peak SeV titers (day 5 PI; both at regular (2×10^5^ pfu) and high dose inoculation) are reduced in atopic mice compared to NA mice. n=3 per group, *p<0.05, ***p<0.001.

### CD11c^+^ cells from atopic mice delay mortality to SeV

We have shown that in atopic mice protection from post-viral airway disease depends upon a PMN dependent process related to increased viral uptake by phagocytic PMNs (Hussain et al., 2021). However, the atopy-dependent protection against mortality did not appear to be due to PMNs, as depletion of PMNs in atopic mice kept all the mice alive and only caused weight loss similar to atopic mice to the viral insult (Supplemental Figure 1A). Similarly, atopic mice deficient in the heme protein myeloperoxidase gene (*Mpo*; important in PMN function and required for extracellular trap formation) were protected from SeV lethality (Supplemental Figure 1B). Since PMNs did not appear to be responsible for the protection against mortality, we examined other cell types. In atopic mice the frequency and numbers of CD11c^+^ cells (dendritic cells or macrophages) were significantly elevated in the airways (Figure 2A). Since we previously demonstrated a critical role for CD11c^+^ cells in the immune axis linking SeV to post-viral airway disease (Hussain et al., 2021; Grayson et al., 2007a; Grayson et al., 2007b), we posited that some of the protection against lethal SeV infection could be due to CD11c^+^ cells from atopic mice. We isolated CD11c^+^ cells from atopic mouse lung and adoptively transferred them into naïve mice. Transfer of these cells 24 h before inoculation with high dose SeV led to a significant delay in mortality, but did not prevent death (Figure 2B). These data suggested that CD11c^+^ cells in atopic mice could be important in providing the survival advantage. To better understand a possible role for CD11c^+^ cells, we analyzed the trancriptional changes in CD11c^+^ cells that occur as a result of making mice atopic. CD11c^+^ cells were isolated from atopic and NA mouse lung and RNA-seq performed. While there were several dysregulated genes, one gene product whose expression was increased several-fold in atopic CD11c^+^ cells was neuregulin-1 (NRG1; Figure 2C). NRG1 is a ligand for ErbB receptors and its expression has been shown to be beneficial in coronary heart disease (Odiete et al., 2012; Geissler et al., 2020), but potentially detrimental in hepatitis C viral infection (Stindt et al., 2016). Given the RNA-seq data and rationale, we measured NRG1 protein levels and found them significantly elevated in the lungs and airways of atopic mice, as well as in *ex-vivo* cultured CD11c^+^ cells isolated from atopic mouse lung (Figure 2D).

**Figure. 2.**
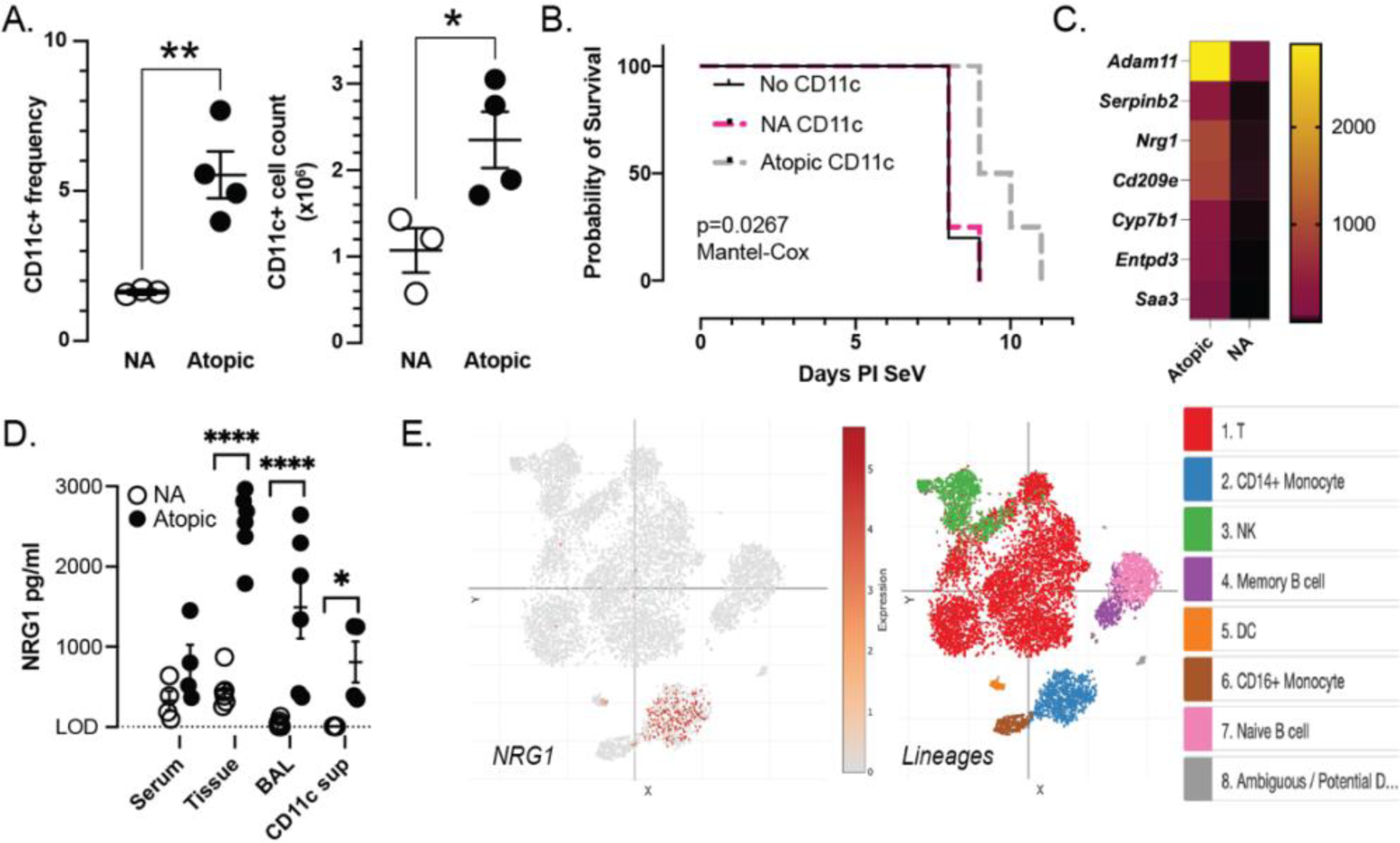
CD11c^+^ cells in atopic mice delay mortality to SeV and produce neuregulin-1 (NRG1). (**A**) CD11c^+^ cells are increased in the lungs of atopic mice. Flow cytometry of lung cell suspension showing frequency (left) and cell number (right) at day 3 post inoculation (PI) high dose SeV (**B**) Adoptive transfer of CD11c^+^ cells isolated from NA or atopic mice into naïve mice 24 h before inoculation with high dose SeV delays but does not prevent mortality; n=4 per group. (**C**) Transcriptomic (RNAseq) comparison between FACS isolated lung CD11c^+^ cells from atopic and NA mice identifies several disparately expressed gene products, including *Nrg1* (n=4 per group). Selected gene products shown. (**D**) NRG1 is markedly increased in atopic mouse lung (“tissue”), BAL, and supernatant from 1 x10^6^ atopic lung CD11c^+^ cells (“CD11c sup”) cultured for 24 h. (**E**) The scRNA (10x Platform) tSNE coordinates show CD14^+^ monocytes and dendritic cell (“DC”) sub-populations expressing NRG1 in peripheral blood cells of healthy human donors.*p<0.05, **p<0.01, ****p<0.0001

Using a publically available database, we found that in healthy human peripheral blood, CD14^+^ monocytes and a subset of dendiritic cells expressed *NRG1* (data source in Methods and Figure 2E). While we have not identified the specific CD11c^+^ cell in the mouse that is making NRG1, these data do suggest that our mouse findings may be applicable to the human.

### NRG1 prevents death from respiratory viral infection *in vivo*

We next assessed the importance of NRG1 in mediating survival from SeV infection. NRG1, a 44-kD glycoprotein, is a cytokine of the epidermal growth factor family and is expressed in 14 isoforms due to alternate splicing or use of alternate promoters. All isoforms contain either an alpha or beta variant epidermal growth factor (EGF)-like domain at their C-terminus that binds to ErbB receptor tyrosine kinases (ErbB1-ErbB4).

The EGF-like motif is sufficient for most biological effects of the full-length ErbB proteins (Ieguchi et al., 2010; Appert-Collin et al., 2015). NRG1 appears to play a homeostatic role with studies showing that loss or overexpression of NRG1 can disrupt NRG1-ErbB signaling (Agarwal et al., 2014; Wang et al., 2021); however, little is known about the role of NRG1 in respiratory viral infections. Given the elevated level of NRG1 in the airways of atopic but not NA mice, we explored whether NRG1 alone could protect mice from respiratory viral induced mortality.

We administered NRG1 (10 to 1000 ng daily) i.n for 5 days to naïve mice before infecting with high dose SeV and determining mortality. In a dose responsive fashion, survival was increased with NRG1 administration (Figure 3A). While NRG1 protected from death, it had a more nuanced effect on post-viral airway disease with mucous cell metaplasia (MCM) increasing, while AHR was blunted but not prevented (Supplemental Figure 2A and B); these data support the contention that the survival mechanism is distinct from that of post-viral airway disease.

**Figure 3.**
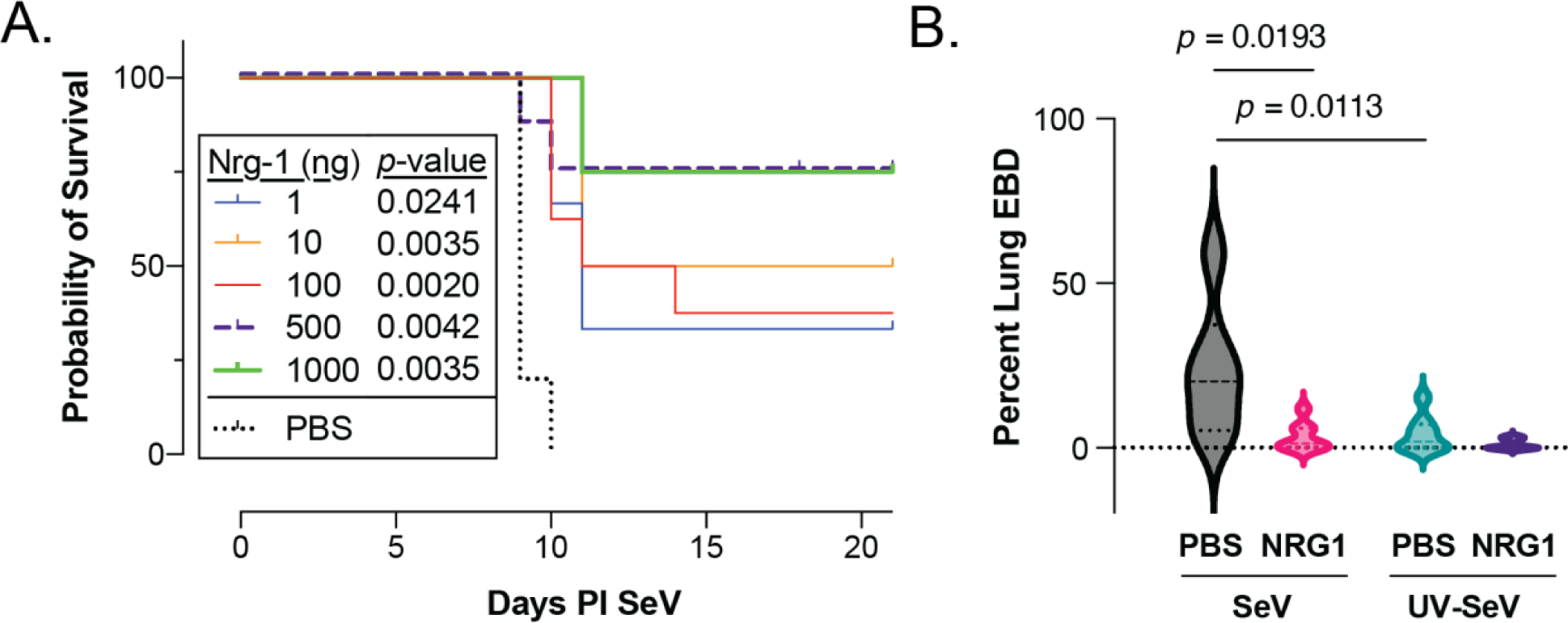
NRG1 is sufficient to reduce SeV induced mortality and airway fluid leak. (**A**) NRG1(1ng to 1000ng) i.n. (in 30µL) given daily to naïve mice for 5d before inoculation with high dose SeV reduces viral mortality; n=4 per group (1ng, 10ng 1000ng & PBS), n=8 per group (100 ng and 500 ng). (**B**) Ratio of EBD in the BAL to that in the lung shows reduced EBD in NRG1 treated mice on day 8 PI SeV. n≥8 per group, median ± IQR shown, Mann-Whitney U test.

It has been reported that pulmonary microvasculature and alveolar epithelium form a barrier function that maintains lung fluid balance. Injury to lung can disrupt this barrier function leading to further exacerbation of lung injury (Mehta et al., 2004; Smith et al., 2021). We hypothesized that one factor leading to death from a severe respiratory viral infection might be both an increase in lung endothelial vascular leakage and epithelial barrier permeability. If this were the case, we would expect NRG1 to reduce leakage during SeV infection. To test this hypothesis, we administered 500 ng NRG1 or PBS for 5 days before infection with SeV (2×10^5^ pfu, the dose at which all mice survive SeV, so that the PBS treated animals would still be alive) or UV-SeV as control and administered Evans Blue dye (EBD) i.v. on day 8 post infection to measure vascular leak. As we hypothesized, there was an increase in lung vascular leak (increased EBD in the lung) and epithelial permeability (as evidenced by EBD in the bronchoalveolar lavage, BAL) with SeV infection in the PBS treated SeV infected animals (Supplemental Figure 3A and B). However, administration of NRG1 prior to viral infection significantly reduced EBD in the BAL, but not lung, suggesting improved epithelial membrane integrity, with no effect on vascular leak (Figure 3B and Supplemental Figure 3A and B).

As noted, we selected a non-lethal SeV dose as on day 8 post infection there would have been significant mortality with high dose SeV (2×10^6^ pfu) in the PBS treated animals. Further studies will be needed to clearly understand the role of NRG1 in contributing to prevention of fluid leak into the airways.

### NRG1 prevents respiratory viral replication *in vitro*

To determine if NRG1 had a direct effect on respiratory viral replication, we utilized *in vitro* human and mouse airway epithelial cell culture systems. The airway epithelium provides the first line of defense against inhaled bacteria and viruses. Therefore, maintenance and integrity of the bronchial epithelium is critical for its barrier function, and our *in vivo* data suggested a role for NRG1 in mediating epithelial integrity. Well-differentiated human bronchial epithelial cell cultures (hBEC) were treated on the baso-lateral side with NRG1 (10 ng, 50 ng, 100 ng) in 500μl media 5 and 3 days before, as well as with, a viral inoculum of 4000 pfu RSV encoding GFP (rgRSV). As can be seen in figure 4 (A and B), RSV replication was reduced (i.e., reduced GFP expression) in NRG1 treated hBEC. A similar result was obtained in mouse tracheal epithelial cultures (mTEC) inoculated with GFP-expressing SeV (GFP-SeV) (Figure 4, A and C).

**Figure 4.**
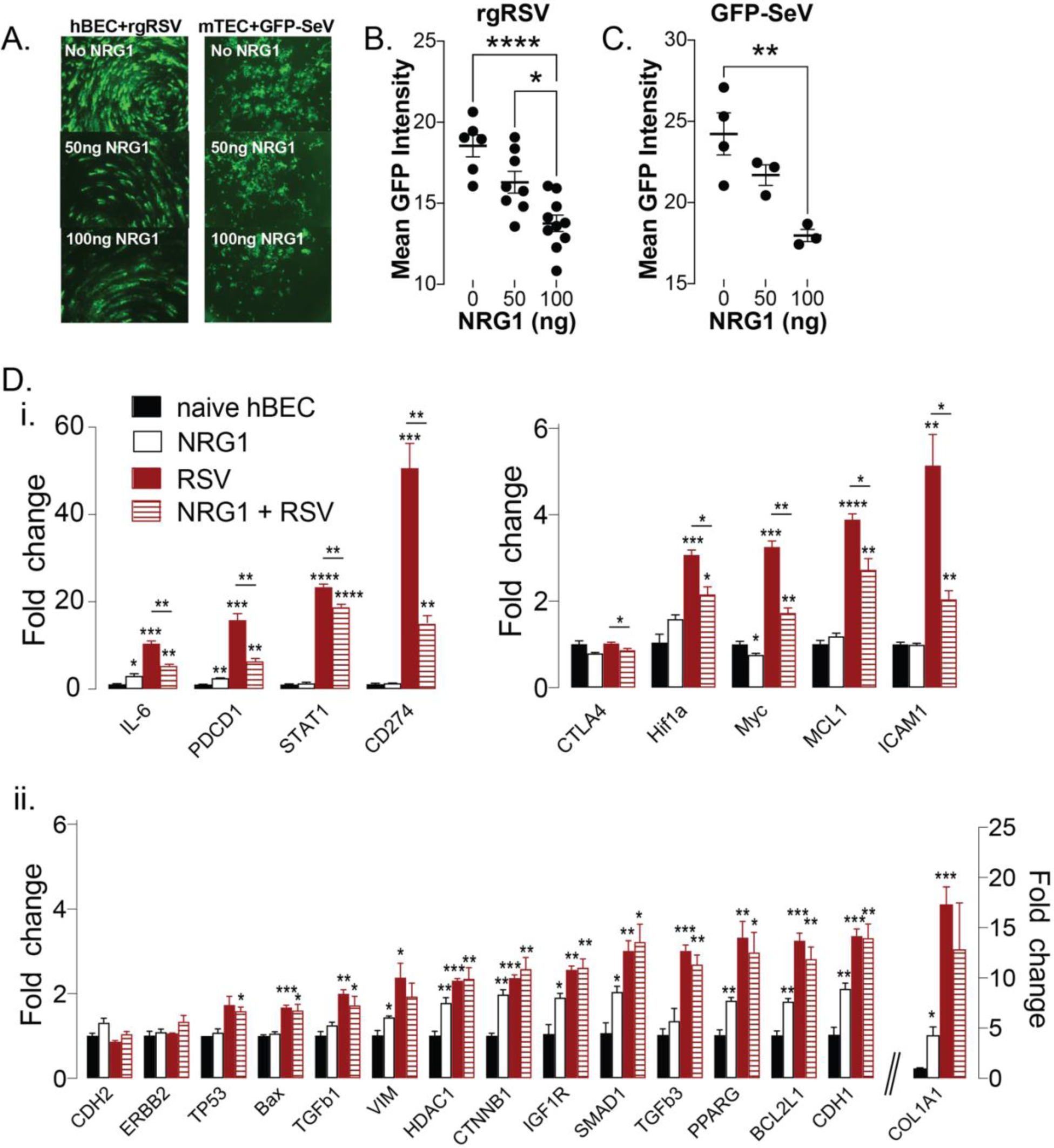
NRG1 treatment reduces viral replication and regulates gene expression in airway epithelium. (**A**) Adding NRG1 to hBEC inoculated with rgRSV (left) and mTEC inoculated with GFP-SeV (right) reduces spread of infection. Representative images shown. (**B**) Quantification of (A) for rgRSV and hBEC and (**C**) for GFP-SeV and mTEC. GFP positive cells quantified by ImageJ. Representative images from ≥3 separate experiments. *p<0.05, **p<0.01. (**D**) Transcriptomic analysis of hBEC cultures treated with NRG1 (100 ng) on the basolateral side of the Transwell for 5 days and inoculated with RSV (4000 pfu). RNA was isolated 48 h PI RSV and qRT-PCR performed using a custom Prime PCR array plate: (i) Transcripts in which NRG1 treatment reduced gene expression from that seen in RSV infected cells. (ii) Transcripts with low level expression that show small but significant change in expression relative to naïve control with NRG1 alone or genes significantly increased with RSV but whose expression levels were not affected by NRG1. *p<0.05, **p<0.01, ***p<0.001, ****p<0.0001, n=3.

### NRG1 modulates homeostasis related genes in epithelial cells

Upon infection of airway epithelium, respiratory viruses such as RSV and rhinovirus augment chronic inflammation and delay viral clearance, while others such as COVID-19, lead to severe damage of the epithelial cell barrier due to cell death (Tan et al., 2020). Since NRG1 is required for maintaining epithelial differentiation via ErbB signaling and promotes epithelial cell proliferation and repair (Vermeer et al., 2006; Jarde et al., 2020; Liu and Kern, 2002; Patel et al., 2000; Kettle et al., 2010), we examined the effect of NRG1 on cellular proliferation in RSV-infected hBEC. NRG1 pretreatment led to increased thickness of the epithelial cell layer in RSV-infected hBEC (Supplemental Figure 4), suggesting that similar to other epithelial injury models (Tan et al., 2020) NRG1 may support epithelial cell proliferation and repair processes in viral infections.

It has been reported that NRG1 plays a critical role in maintaining homeostasis potentially by reducing cellular and mitochondrial stress (Zhang et al., 2016; Guma et al., 2020; De Keulenaer et al., 2019). Using a custom gene array and NRG1 treated or untreated RSV-infected hBEC, we evaluated expression of gene transcripts associated with a homeostatic/protective function of airway epithelium. Administration of NRG1 to uninfected epithelial cells had little effect on the expression of most genes examined (Figure 4D). However, infection with RSV led to a marked and significant upregulation in many genes. Importantly, treatment with NRG1 reduced expression of nine genes – *IL6*, *PDCD1*, *STAT1, CD274 (PD-L1), CTLA-4, HIF1a*, *MYC*, *MCL1*, and *ICAM-1* (Figure 4D(i)). Further studies are needed to determine functional relationships between these genes and their protein products.

The expression of *IL6* is interesting, as this cytokine has both pro-and anti-inflammatory functions (Scheller et al., 2011; Tanaka et al., 2014). *IL6* was increased by RSV infection of hBEC; however, NRG1 treatment of epithelial cells significantly attenuated this upregulation of *IL6* expression 48 h post inoculation RSV (Figure 4D(i) left panel) supporting a potential role for NRG1 in regulation of inflammatory responses.

PD-1, encoded by *PDCD1* mitigates inflammatory cell activation and contributes to mitochondrial dysfunction (Yao et al., 2015; Ogando et al., 2019), while its ligand, PD-L1, has increased expression on lung epithelial cells. Thus, potentially by impairing expression of both PD-1 and PD-L1, NRG1 could increase the antiviral inflammatory response leading to increased survival. Supporting this hypothesis is an *in vitro* hBEC study that demonstrated blocking PD-L1 in acute RSV infection facilitated the antiviral immune response (Telcian et al., 2011). Downregulation of the PD-L1-PD1 axis by NRG1 further suggests a protective role in acute respiratory infection.

Some viruses including RSV delay apoptosis of infected cells to prevent premature death of host cells by increasing anti-apoptotic MCL-1 expression to promote viral latency and persistent infection (Schwarze et al., 2004; Kotelkin et al., 2003). Our data supports RSV infection increasing *MCL1* with a significant reduction in *MCL1* expression in hBEC pre-treated with NRG1 prior to RSV infection; however, *MCL1* is still significantly higher in the NRG1+RSV group than in cells treated with NRG1 without RSV infection (Figure 4D(i), right panel). These data suggest that NRG1 may induce controlled (or selective) apoptosis in infected cells as a means of protection from the viral insult. Further, caspase-3, an apoptotic effector caspase, has been shown to cleave MCL-1 thus facilitating apoptosis (Weng et al., 2005). Although we have not looked at the expression of MCL-1 protein in mouse lung epithelium, we did find increased activated caspase-3 at day 5 PI SeV in small airways of NRG1 treated mice (Supplemental Figure 5). The increased levels of active caspase-3 suggest that NRG1 in mice induces a similar pro-apoptotic state as we see in the hBEC.

ICAM-1 has been shown to facilitate RSV entry and infection of epithelial cells (Behera et al., 2001); therefore, reduced expression of *ICAM-1* in NRG1 treated RSV infected hBEC could contribute to reduced virus spreading. Moreover, *ICAM-1* expression is still significantly higher in NRG1 treated and infected hBEC than in untreated or uninfected NRG1 treated epithelial cells suggesting a potential role in tissue repair and resolving inflammation (Figure 4D(i)), (Bui et al., 2020). MYC activation has been shown to provide support for respiratory virus replication including IAV and adenovirus (Thai et al., 2015). Therefore, a reduction in *MYC* expression in our study further suggests NRG1 could be inhibiting RSV replication (Figure 4D(i)). HIF1α, a transcription factor, has been implicated in viral infections vesicular stomatitis virus and herpes simplex virus) and in the cytokine storm seen with severe COVID-19 disease (Tian et al., 2021). Increased *HIF1α* is often a consequence of mitochondrial damage, which will impair epithelial repair similar to what we have reported with mouse *Hif1α* and re-epithelialization of skin wounds (Biswas et al., 2010). Further, HIF1α has been shown to increase epithelial permeability (Song et al., 2017). Therefore, a reduction in *HIF1α* expression in hBEC with NRG1and RSV further supports the notion that NRG1 is a homeostatic regulator of viral infection (Figure 4D(i)). Finally, there were genes with small but significant increases in expression in NRG1 treated hBEC relative to naïve cells (Figure 4D(ii)). Altogether, we posit that these data suggest NRG1 reduces the cellular stress imposed by RSV on infected hBEC, while at the same time increasing apoptosis of these cultured cells.

In conclusion, our studies demonstrate that the presence of atopy before a severe paramyxoviral infection is sufficient to protect against an otherwise lethal infection. This protection appears to be due to increased expression of NRG1, although we acknowledge that the protection seen in HDM treated mice may be due to other mechanisms. Nonetheless, intranasal administration of NRG1 alone is sufficient to protect mice from a lethal viral infection, and administration of NRG1 to epithelial cell cultures *ex vivo* is sufficient to reduce RSV (for human epithelial cells) and SeV (for mouse epithelial cells) replication. This reduced viral titer is associated with a decrease in expression of the PD-1, PD-L1 axis, and *in vivo* correlates with reduced fluid leak into the airways. Together, we have identified a novel role for NRG1 in providing protection from a severe respiratory viral infection. Future studies will explore in more detail the mechanism of NRG1 mediated protection, as well as the therapeutic potential of NRG1 for the treatment of severe respiratory viral infections.

## Materials & Methods

### Ethics statement

All animal studies were performed under the protocol approved by the Institutional Animal Care and Use Committee of the Abigail Wexner Research Institute at Nationwide Children’s Hospital. Human tissues for cell culture were obtained from de-identified donors. The progenitor cells for these cultures were extracted from de-identified lung tissue provided by The Ohio State University Wexner Medical Center, Comprehensive Transplant Center Human Tissue Biorepository and is exempted from IRB approval.

### Animals

C57BL/6 (wild type) mice and B6.129X1-*Mpo*^tm1lus/J^ (common name: MPO KO (strain#: 004465)) were purchased from the Jackson Laboratory (Bar Harbor, Maine, USA) and bred in-house. Unless indicated otherwise mice 8-12 weeks old were used for the experiments.

### Human and mouse air-liquid interphase culture

Human bronchial epithelial culture (hBEC) progenitors and C57BL6 mouse tracheal epithelial cultures (mTEC) were grown on transwells at the air-liquid interface for 5 to 6 weeks for the development of pseudostratified well-differentiated airway epithelial layers resembling *in vivo* epithelium with ciliated, goblet and basal cell types as we and others have published (Fulcher et al., 2005; King et al., 2021; Eenjes et al., 2018; Hussain et al., 2021).

### Atopic model and SeV infection

C57BL6 mice were sensitized with 1 μg for one day and challenged for five days with 10 μg of house dust mite (HDM) extract (catalog no. XPB91D3A2.5; Stallergenes Greer USA, Boston, MA) given intranasally (i.n.) to make them atopic or with PBS as non-atopic (NA) control. Three days after last challenge mice were inoculated i.n. with 2×10^5^ pfu (regular dose) or 2×10^6^ (high dose) SeV (Hussain et al., 2021).

### Flow cytometry and flow-activated cell sorting (FACS) of CD11c^+^ cells

Cells from whole mouse lungs were harvested and flow cytometry and FACS performed as we previously reported using standard cell staining techniques with CD11c antibody (Clone N418; catalog no. 12-0114-82, ThermoFisher) or Armenian Hamster IgG Isotype Control (Clone: eBio299Arm, catalog no. 12-4888-81, ThermoFisher), (Cheung et al., 2010a; Grayson et al., 2018; Hussain et al., 2021).

### Adoptive transfer and SeV infection

Lung CD11c^+^ cells purified (≥85% CD11c^+^) from NA and atopic mice by FACS were delivered intranasally to anesthetized naïve C57BL6 mice at a concentration of 1.0 x 10^6^ cells/mouse. After 24 h recipient mice were inoculated with 2×10^6^ pfu SeV and survival recorded over days post inoculation.

### Airway hyper-reactivity (AHR) and mucous cell measurement

Invasive measurement of AHR was performed by measuring airway resistance to increasing doses of methacholine using the FlexiVent system as we have published (Cheung et al., 2010b; Hussain et al., 2021). Mucous cell metaplasia was determine by performing Periodic-Acid-Schiff (PAS) staining of formalin fixed lung sections and in a blinded fashion, manually counting and determining the number of PAS^+^ cells per mm of basement membrane (using ImageJ) as we have reported (Hussain et al., 2021).

### Immunohistochemistry

Paraffin embedded mouse lung tissues sectioned at 10μM thickness were dewaxed followed by heat induced antigen retrieval and stained with active caspase-3 antibody (catalog #: AF835-SP, R & D Systems) at 1μg/ml overnight at 4°C followed by incubation with the Anti-Rabbit IgG VisUCyte™ HRP Polymer Antibody (catalog # VC003, R & D Systems). Tissues were stained with DAB (brown) and counterstained with hematoxylin (blue) following manufacturer’s instructions.

### RNA sequencing

Whole transcriptome profiling was performed by preparing strand-specific RNA-seq libraries using NEB Next Ultra II Directional RNA Library Prep Kit for Illumina, following the manufacturer’s recommendations. In summary, total RNA was assessed using RNA 6000 Nano kit on Agilent 2100 Bioanalyzer (Agilent Biotechnologies) and Qubit RNA HS assay kit (Life Technologies). A 140-500 ng aliquot of total RNA was rRNA depleted using NEB’s Human/Mouse/Rat RNAse-H based Depletion kit (New England BioLabs). Following rRNA removal, mRNA was fragmented and then used for first-and second-strand cDNA synthesis with random hexamer primers and ds cDNA fragments undergoing end-repair and a-tailing and ligation to dual-unique adapters (Integrated DNA Technologies). Adaptor-ligated cDNA was amplified by limit-cycle PCR. Library quality was analyzed on Tapestation High-Sensitivity D1000 ScreenTape (Agilent Biotechnologies) and quantified by KAPA qPCR (KAPA BioSystems). Libraries were pooled and sequenced at 2 x 150 bp read lengths on the Illumina HiSeq 4000 platform to generate approximately 60-80 million paired-end reads per sample. Differential expression analysis was performed and significant differentially expressed features between the two groups with an absolute value of fold change ≥ 1.5 and an adjusted p-value of ≤ 0.10 (10% FDR) were recorded.

### NRG1 ELISA

Detection of NRG1 levels was performed using mouse NRG1 ELISA kit (cat# EKN47308, Biomatik, Delaware USA) following manufacturer’s instructions. Freshly prepared standards, tissue extract and cell supernatant were used for the assay and results quantified by plotting against the standard curve with detection range of 15.6-1000 pg/ml and reading O.D. at 450 nm.

### NRG1 exogenous administration and cell culture treatment

Recombinant mouse neuregulin-1/NRG1 protein, CF (cat#: 9875-NR; Novus Biologicals, CO, USA) of varying concentrations (10 to 500 ng) were given i.n. daily for 5 days before inoculating with SeV. Data were recorded for weight change and survival for two weeks post infection. Cell culture basal media was supplemented with recombinant human NRG1 alpha (cat#: NBP2-35093, Novus Bio) for hBEC or with mouse NRG1 for mTEC on day five and three before and at the time of infection with 4000 pfu of rgRSV or SeV-GFP. Virus inoculation occurred in 100μl DMEM on apical side for 4 hrs at 37°C in 5% CO_2_ incubator. GFP levels were quantified by imaging with EVOS cell imaging system at 10x magnification and determining the mean fluorescent intensity with ImageJ software (Schneider et al., 2012). Cell culture harvested after 48 h for RNA isolation and qRT-PCR. The SeV-GFP was a gift from Dr. Dominique Garcin, Switzerland and the construct is published (Strahle et al., 2007) and rgRSV, that also encodes GFP, was developed by our group (Kwilas et al., 2010).

### Vascular leak and epithelial permeability quantification

Vascular leak and alveolar permeability were assessed by measuring accumulation of Evans blue dye (EBD) in the lung or bronchoalveolar lavage (BAL) respectively with modifications as described by our group and others (Carpenter and Stenmark, 2000; Smith et al., 2021; Kelly et al., 2016). Briefly, 100 µL EBD (Sigma-Aldrich cat# E2129) at 20 mg/kg was injected i.v. via the tail vain. One hour later bronchoalveolar lavage was performed, BAL fluid collected and lungs were harvested, then formamide was added to each sample to extract EBD. The extracted dye was quantitated by spectrophotometry, measuring absorbance at 620 nm with absorbance at 740 nm used as a baseline control.

### RNA Isolation and Quantitative Real-Time PCR

Total RNA from hBEC isolated using mirVana miRNA and total RNA isolation kit (catalog no. AM1560, Thermo Fisher Scientific). For qRT-PCR cDNA was synthesized using Maxima H Minus cDNA synthesis kit (catalog no. M1681; Thermo Fisher Scientific). PrimePCR^TM^ PCR Primers assay plates were custom made with validated primers for use with EvaGreen dye-based chemistry (Bio-RAD, USA). Samples were run on CFX96 Touch Real-Time Detection System (Bio-RAD, USA). C_T_ value normalized with *GAPDH* or *MRPL13* and expressed as fold change relative to naïve control. For the array we selected 24 genes that broadly fall into three groups: (i) epithelial to mesenchymal transition signature genes; (ii) genes in RNA virus infection panel of pre-designed human PrimePCR by BioRad; (iii) fibroblast genes previously reported to be regulated by NRG1 (De Keulenaer et al., 2019).

The following human genes were included in the assay: *BAX* (BCL2-associated X protein); *BCL2L1* (BCL2-like 1); *CD274* (PD-L1); *CDH1* (cadherin 1, type 1, E-cadherin (epithelial); *CDH2* (cadherin 2, type 1, N-cadherin (neuronal); *COL1A1* (collagen, type I, alpha 1); *CTLA4* (ICOS, cytotoxic T-lymphocyte-associated protein 4); *CTNNB1* (catenin (cadherin-associated protein), beta 1, 88kDa), *GAPDH* (glyceraldehyde-3-phosphate dehydrogenase); *HDAC1* (histone deacetylase 1); *HIF1A* (hypoxia inducible factor 1); *ICAM1* (intercellular adhesion molecule 1); *IGF1R* (insulin-like growth factor 1 receptor); *IL6* (interleukin 6); *MCL1* (myeloid cell leukemia sequence 1 (BCL2-related)); *MRPL13* (mitochondrial ribosomal protein L13); *MYC* (v-myc myelocytomatosis viral oncogene homolog (avian); *PDCD1* (programmed cell death 1); *PPARG* (peroxisome proliferator-activated receptor gamma); *SMAD1* (SMAD family member 1); *STAT1* (signal transducer and activator of transcription 1, 91kDa); *TGFB1* (transforming growth factor, beta 1); *TGFB3* (transforming growth factor, beta 3); *TP53* (tumor protein p53); *VIM* (vimentin).

### SingleCell RNAseq Data source

Immune Cell Atlas: Blood Mononuclear Cells, Single Cell Portal from Broad Institute was used to derive NRG1 expression data from healthy human PBMC profiled by SingleCell RNAseq (10x platform). https://singlecell.broadinstitute.org/single_cell/study/SCP345/ica-blood-mononuclear-cells-2donors-2-sites?genes=NRG1#study-summary (Accessed June 8, 2022

### Statistical Analysis

All statistical analyses were performed using Prism 9 (GraphPad Software Inc.). All normally distributed data are presented as mean ± SEM with survival analyses using Kaplan-Meier curves and log-rank (Mantel-Cox) tests; non-normally distributed data are shown as median ± inter-quartile range (IQR). Student’s *t* test (for normally distributed data) or Mann-Whitney U (for non-normally distributed data) was used to assess significant differences between two means. For all tests, \**p*≤0.05; \*\**p*≤0.01, \*\*\**p*<0.001; \*\*\*\**p*≤0.0001 was considered statistically significant.

## Acknowledgments

This work was supported by NIH grants R01HL087778; R01AI171027 (to MHG); R01 AI093848 and U19AI42733 (to MEP). Robert & Edgar Wolfe Foundation and The Abigail Wexner Research Institute (AWRI) at Nationwide Children’s Hospital (to MHG).

The authors thank the Genomic Services at AWRI for performing RNAseq and Cure Cystic Fibrosis Columbus (C3) Epithelial Cell Core (ECC) at Nationwide Children’s Hospital and The Ohio State University for providing primary human bronchial cultures, advice, and tools for this work. C3 is supported by a Research Development Program Grant (MCCOY17R2) from the Cystic Fibrosis Foundation. The source tissue for these cultures was provided by the Comprehensive Transplant Center Human Tissue Biorepository of The Ohio State University Wexner Medical Center or by Nationwide Children’s Hospital.

