## Supplemental Figures 1-5 for "Neuregulin-1 protects against respiratory viral induced mortality"

Hussain et al., 2023

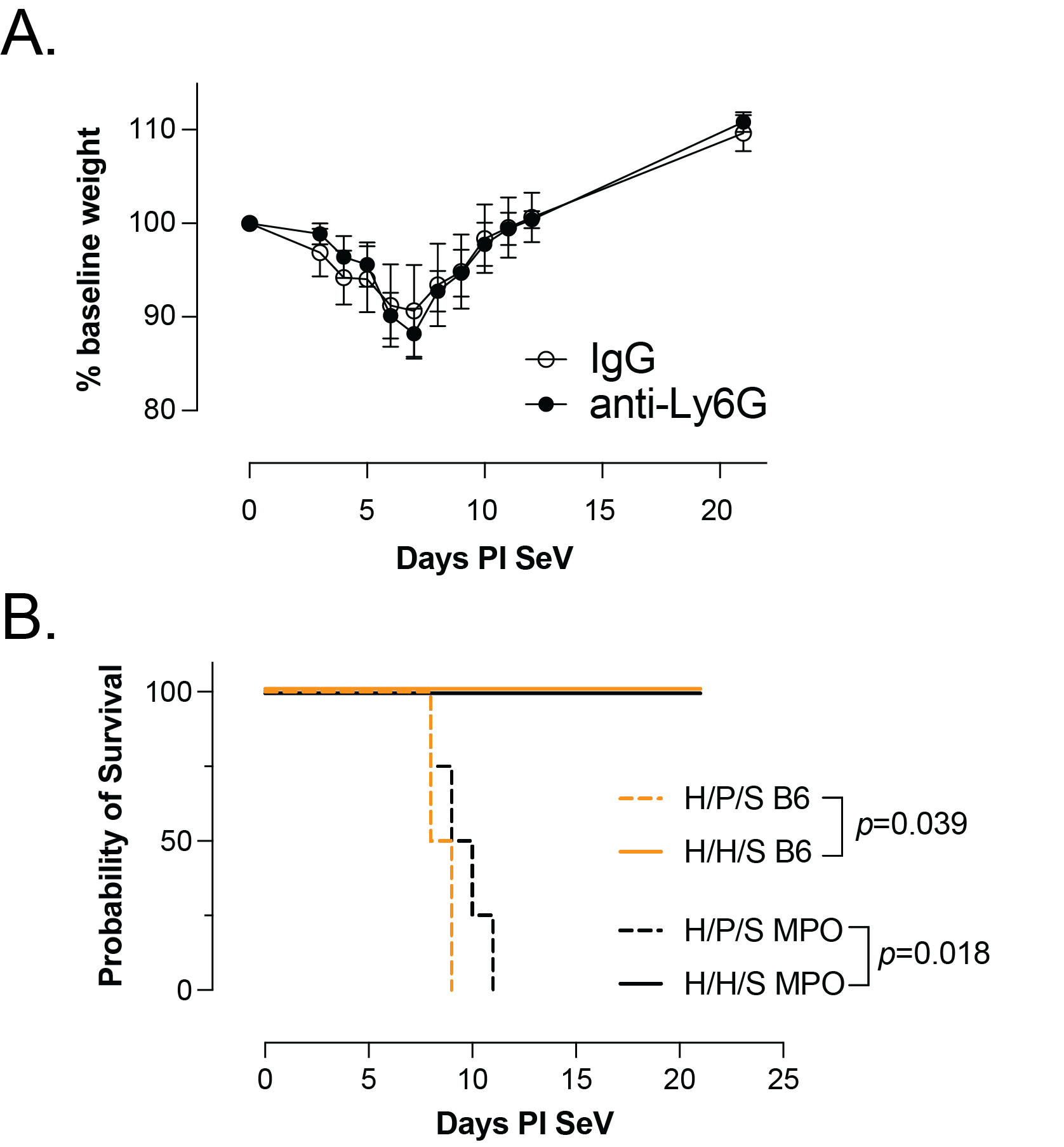

**Supplemental Figure S1: Protection against severe SeV infection is neutrophil independent.** (**A**) Weight loss in atopic mice treated with anti-Ly6G or control IgG 24 h before inoculation with high dose SeV. All mice survived regardless of treatment (n=4). (**B**) Survival curve of atopic (“H/H/S”) or NA (“H/P/S”) wild-type (“B6”) or *Mpo^–/–^* (“MPO”) mice inoculated with high dose SeV; n=3 mice per group.

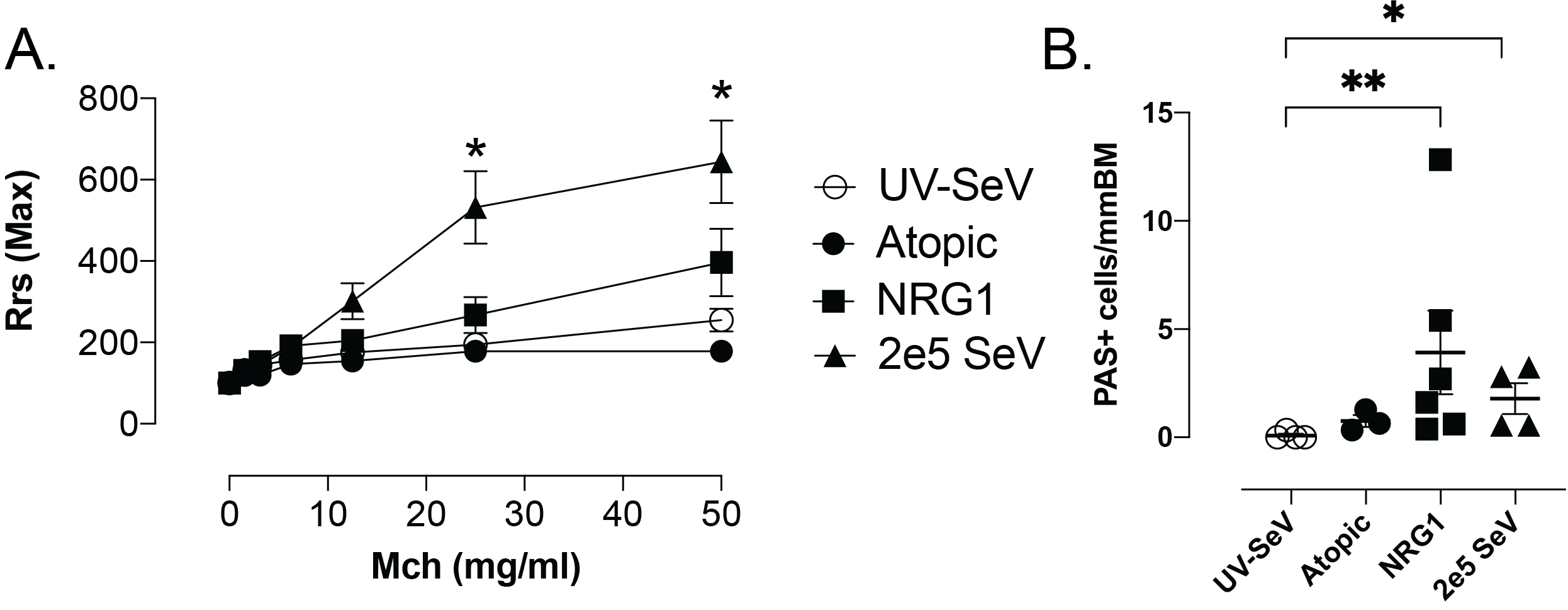

**Supplemental Figure S2:** **NRG1 partially reduces post-viral airway disease**. NRG1 (500 ng) delivered i.n. for 5 days (d-4 to d0; “NRG1”) before inoculation with high dose (2x10^6^ pfu) SeV (**A**) led to a slight (but not significant) increase in airway hyper-reactivity but did significantly increase (**B**) PAS^+^ cells (mucous cell metaplasia) 21 d post infection when compared to ultraviolet light inactivated SeV (UV-SeV) controls. N≥3 per group; for comparison “2e5 SeV” group (regular dose SeV (2x10^5^ pfu) in NA mice) and atopic mice with 2x10^6^ pfu SeV are shown.

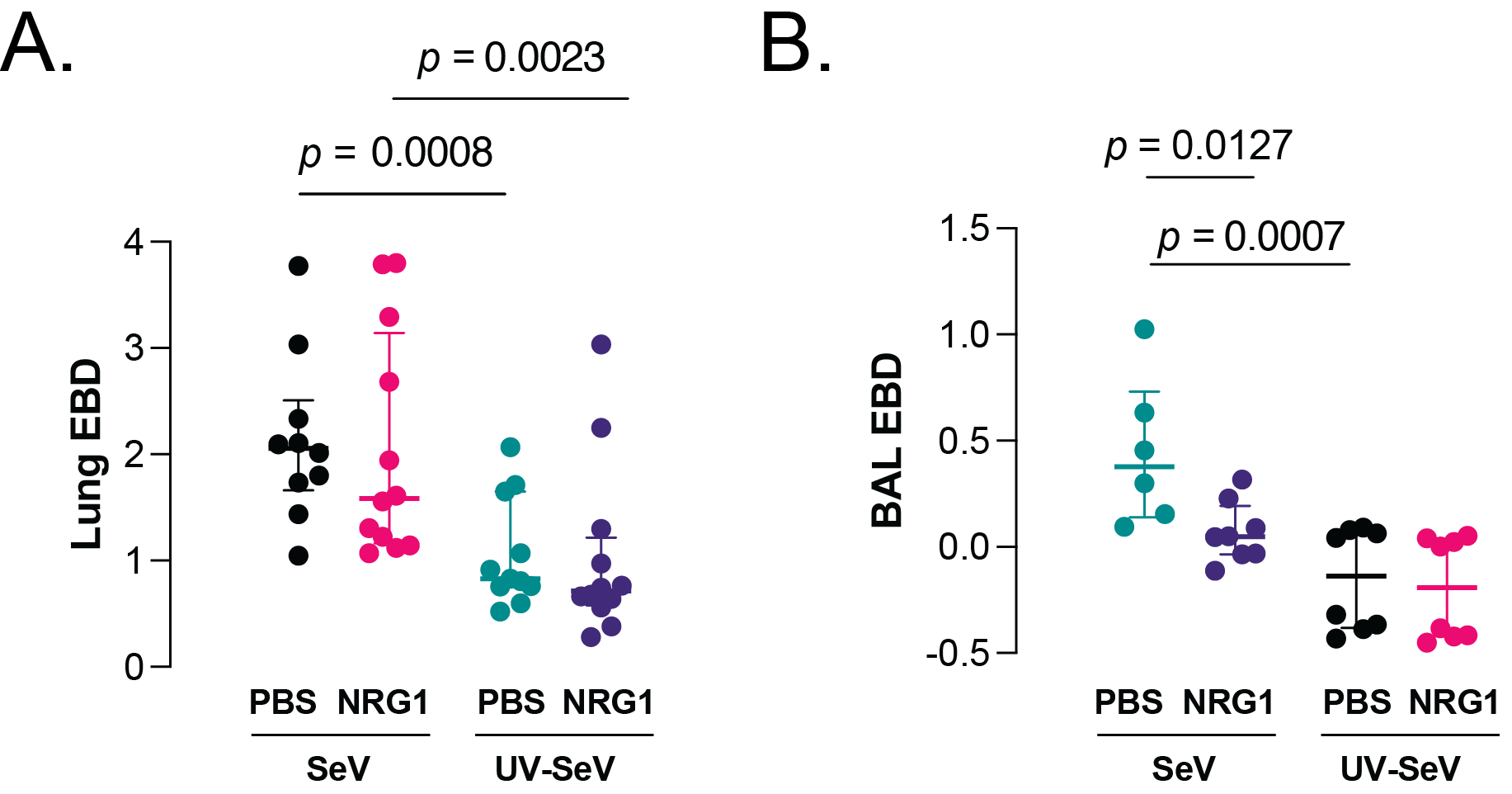

**Supplemental Figure S3: Lung vascular and epithelial permeability. (A**) Lung (n≥10) and (**B**) BAL (n≥6) EBD extravasation at day 8 post inoculation SeV or UV-SeV. Note that NRG1 treatment significantly reduced the amount of EBD in the BAL with SeV infection but had no effect on lung levels.

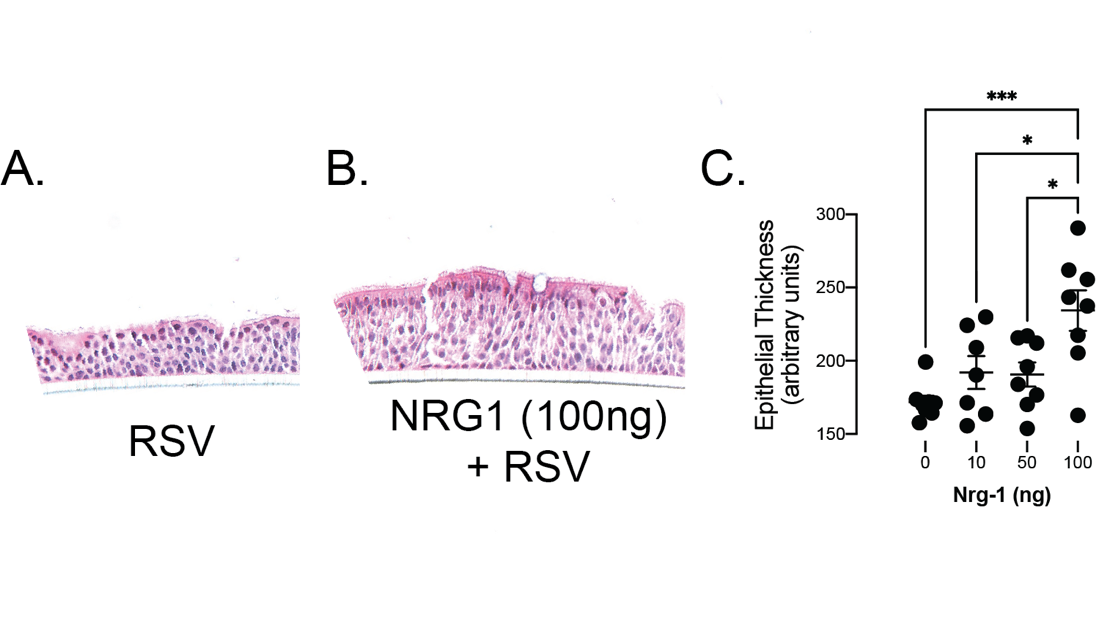

**Supplemental Figure S4. NRG1 increases hBEC thickness.**

Representative H & E staining (20x) of paraffin fixed sections of hBEC cultures 48 h after inoculation with (**A**) RSV (4000 pfu) or (**B**) NRG1 (100 ng) and RSV. (**C**) Quantification of (A) and (B). Epithelial thickness measured with ImageJ software. n≥4 per group.

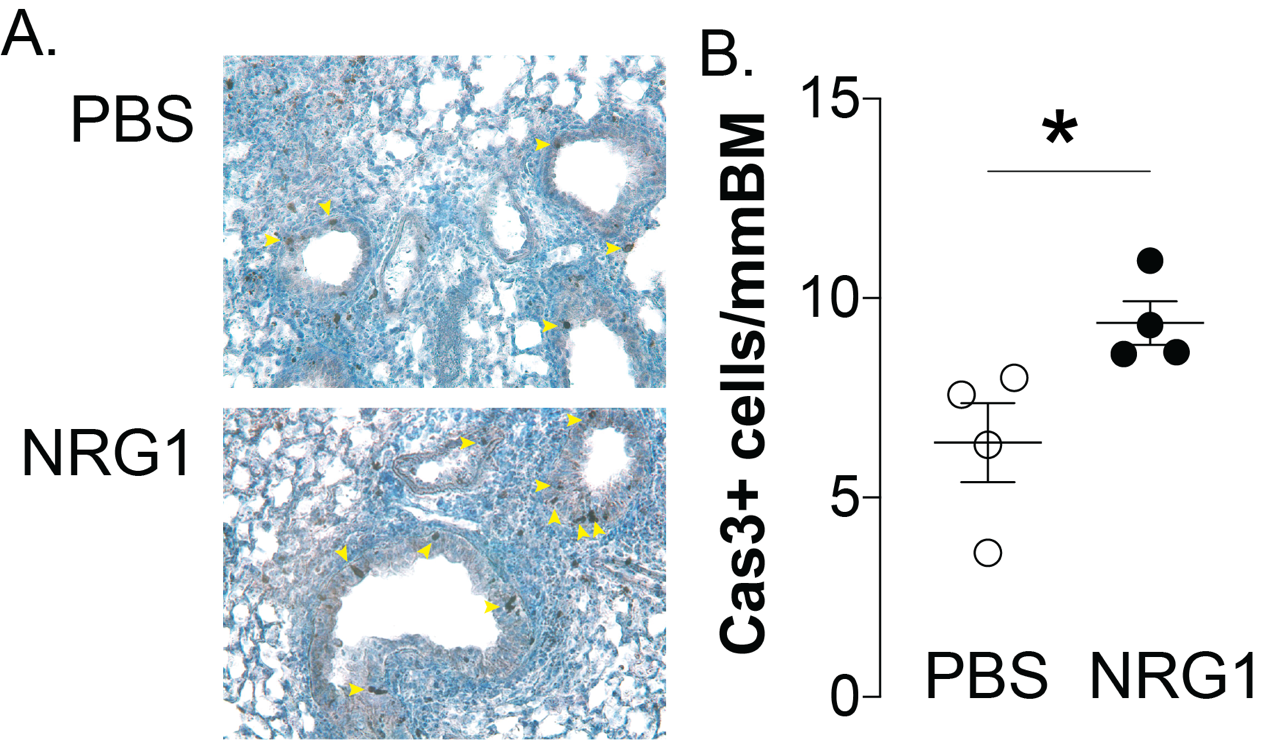

**Supplemental Figure S5: Active caspase-3 increased in small airways of NRG1 treated and virally infected mice. (A)** Representative images (at 20x magnification) of immunohistochemistry of lungs showing active caspase-3 stained epithelial cells (arrow heads) in PBS (top panel) and NRG1 treated (bottom panel) (500 ng, daily day-4 to day 0) at day 5 post inoculation SeV. (**B**) Quantification of (A) expressed per mmBM. *p<0.05
